# Genome-scale genetic interactions position the Mitotic Exit Network as a major antagonist of transient Topoisomerase II deficiency

**DOI:** 10.1101/108761

**Authors:** Ramos-Pérez Cristina, Grant W Brown, Machín Félix

## Abstract

Topoisomerase II (Top2) is the essential protein that resolves DNA catenations. When the Top2 is inactivated, mitotic catastrophe results from massive entanglement of chromosomes. Top2 is also the target of many first-line anticancer drugs, the so-called Top2 poisons. Often, tumours become resistant to these drugs by downregulating Top2. Here, we have compared two isogenic yeast strains carrying *top2* thermosensitive alleles that differ in their resistance to Top2 poisons, the broadly-used poison-sensitive *top2-4* and the poison-resistant *top2-5*. We found that *top2-5* transits through anaphase faster than *top2-4*. In order to define the biological importance of this difference, we performed genome-scale Synthetic Gene Array (SGA) analyses during chronic sublethal Top2 downregulation and acute, yet transient, Top2 inactivation. We find that downregulation of cell cycle progression, especially the Mitotic Exit Network (MEN), protects against Top2 deficiency. In all conditions, genetic protection was stronger in *top2-5*, and this correlated with destabilization of anaphase bridges by execution of MEN. We suggest that mitotic exit may be a therapeutic target to hypersensitize cancer cells carrying downregulating mutations in *TOP2*.

## INTRODUCTION

Upon chromosome replication, topological intertwining arises between sister chromatids. These intertwines often become interlocked (i.e. catenations) due to the confinement of the very long chromosomes in the reduced space of the nucleus. Catenations preclude sister chromatid segregation at anaphase, and the key enzyme in all life forms for removing them is toposimerase II (Top2) (Nitiss 2009a; Vos et al. 2011). Top2 works by making transient double-strand breaks (DSBs) on one sister chromatid, allowing the passage of the sister through it. Importantly, a human homologue of Top2, hTOPOIIα, is the main target of first-line anticancer drugs including etoposide and doxorubicin (Deweese and Osheroff 2009; Nitiss 2009b). These drugs trap Top2-mediated DSBs and are so called Top2 poisons. The resulting DSBs are more abundant and less efficiently repaired in cancer cells than in normal cells and this, in turn, leads to the selective killing of the tumour. The hTOPOIIα is often mutated and/or downregulated during acquisition of secondary resistance to Top2 poisons, and this fact could be exploited for second-line anticancer treatments (Nitiss 2009b; Holohan et al. 2013; Larsen, Escargueil, and Skladanowski 2003).

Top2 is essential for cellular viability. In unicellular eukaryotes and bacteria the study of Top2 functions has been largely facilitated by the availability of conditional alleles. In the yeasts *Saccharomyces cerevisiae* and *Schizosaccharomyces pombe* early studies showed that inactivation of Top2 by means of thermosensitive (ts) alleles leads to a mitotic catastrophe as determined by a sudden loss of viability once the cells reach anaphase (Uemura and Tanagida 1986; Holm et al. 1985). In agreement with a role in removing sister chromatid catenations, Top2 inactivation yielded cells with DAPI-stained anaphase bridges and broken chromosomes once cells completed cytokinesis (Uemura and Tanagida 1986; Holm et al.1985; Uemura and Yanagida 1984; DiNardo, Voelkel, and Sternglanz 1984; Stearns, and Botstein 1989). In the case of *S. cerevisiae*, all these studies were carried out with two ts alleles isolated in independent screens, *top2-1* and *top2-4*. Both alleles yield Top2-ts proteins sensitive to poisons. In the same screen where *top2-4* was isolated, *top2-5* was also obtained (Holm et al. 1985). Later, *top2-5* was shown to be resistant to poisons and served as a key tool to understand the mechanism of action of this class of clinical drugs (Jannatipour, Liu, and Nitiss 1993; Perego et al. 2000). Nevertheless, the *top2-5* cell cycle was not characterized and has been assumed to be equivalent to that of *top2-4*.

Here, we have revisited the cell cycle progression of cells expressing the broadly used *top2-4* allele and compared its behaviour to an isogenic *top2-5* strain. In addition, we have performed a genome-scale synthetic genetic array (SGA) analysis for these two *top2-ts* alleles. We show that execution of the Mitotic Exit Network has specific deleterious effects on sublethal downregulation of Top2-5.

## RESULTS AND DISCUSSION

### Cells carrying the *top2-5* allele complete anaphase faster than *top2-4*

We started this work by revisiting fluorescence microscopy time course experiments in the original *top2-4* and *top2-5* ts strains (Holm et al. 1985). As a control, we also included the *TOP2* reference wild type allele in the same genetic background. We first engineered the strains in order to label the histone H2A (*HTA2* gene) with GFP. This strategy allowed us to complement the time course experiments with fluorescence videomicroscopy of the nuclear DNA without adding DNA intercalating dyes. We further labelled Rad52 with RedStar2 in order to detect whether cells suffered DNA damage upon inactivation of Top2. Rad52 forms nuclear foci to repair DSBs through the homologous recombination repair pathway (Lisby, Rothstein, and Mortensen 2001).

All strains were arrested in G1 at the permissive temperature (25 ºC) for 3 hours and then released at 37 ºC to follow the progression through a synchronous cell cycle with Top2-ts inactivated. As expected, the *TOP2* control cycled normally after the release, with normal segregation of the nuclear masses (i.e., long-distance binucleated, LDB) taking place very quickly at 90’-120’ and no signs of DNA damage after that (Figure 1a, left panels). By contrast, the *top2-4* mutant exhibited a phenotype of cells stuck in anaphase/telophase after 4 hours, in which approximately 70% of the cells were in a “dumbbell” state (i.e., the bud as big as the mother) and about 35% had two very close split nuclear masses (i.e., short-distance binucleated, SDB) as determined by histone-labelling (Figure 1a, central panels). The presence of this abnormal form of nuclear segregation was constant from 150’ onward. Interestingly, the same SDB phenotype was described before for a strain that depletes Top2 through a degron system and referred as “cut” phenotype (Baxter and Diffley 2008). Besides, a similar terminal phenotype occurs in mouse topo IIα^−/−^ embryos and in a significant fraction (~1/3) of epithelial cells treated with Top2 catalytic inhibitors (Akimitsu et al. 2003; Wheatley, O’Connell, and Wang 1998). Coinciding with this change in the nuclear morphology, a steady increase of cells with Rad52 foci was also observed (up to 35% of cells by 240’). Strikingly, *top2-5* had a very different mix of cell morphologies, which included a decrease of the dumbbell category from 150’ and the presence of “threesomes”, where the mother cell has re-budded, in up to 25% of the population (Figure 1a, right panels). Furthermore, there was only a transient peak of SDB at 120’, with less than 10% of the cells having the SDB morphology after 4 hours. Likewise, Rad52 foci abruptly rose from 120’, reaching 60% of all cells by 180’ (twice as many as in *top2-4*).

**Figure 1.**
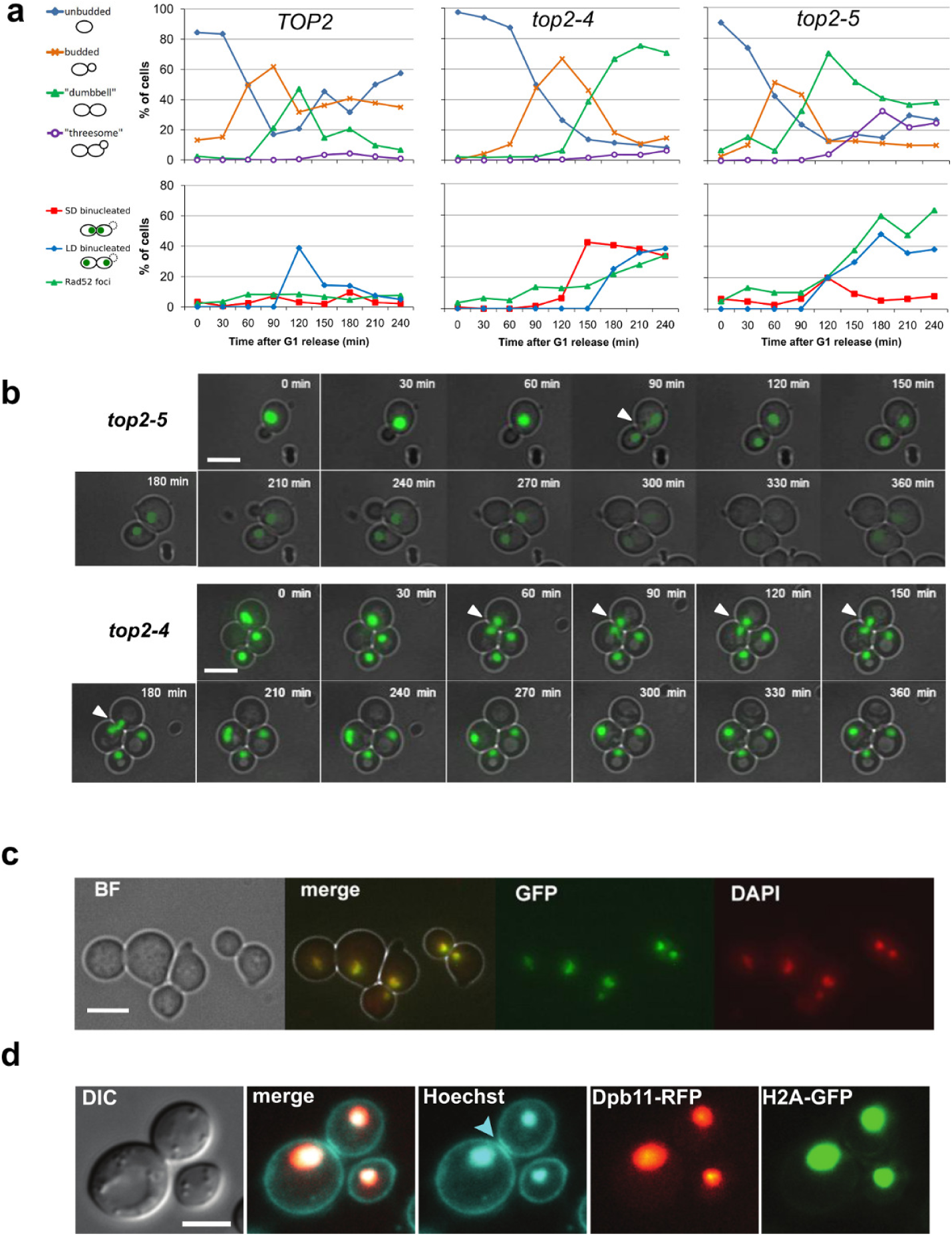
Anaphase bridges in *top2-5* are quickly split apart at anaphase. **(a)** *HTA2-GFP Rad52-RedStar2* yeast cells carrying different alleles of the *TOP2* gene (wild type *TOP2* and thermosensitive alleles *top2-4* and *top2-5*) were synchronized in G1 at the permissive temperature (25ºC) and then released into a synchronous cell cycle at 37ºC for 4 h. Samples were taken every 30′ and directly analysed by fluorescence microscopy. Upper charts depict the budding pattern of the population. Lower charts depict the percentage of cells with histone-labelled split nuclear masses that either remained at a short distance (≤2 µm) from one another (“SD binucleated”) or were separated by a longer distance (“LD binucleated”). Percentage of cells with at least one Rad52 focus is also included. **(b)** *HTA2-GFP* cells carrying the *top2-ts* alleles were grown at 25ºC, concentrated to OD_600_=3, spread onto YPD agarose patches and filmed under the microscope at 37ºC for 6 hours, taking images every 30’. Representative cells are shown. White filled triangles point to chromatin bridges visible with H2A-GFP. **(c)**
*HTA2-GFP top2-4* after 4 hour at 37ºC in liquid cultures showing a perfect colocalization of H2A-GFP and DAPI staining. **(d)**
*HTA2-GFP Dpb11-yEmRFP top2-5* cells were grown to exponential phase in synthetic complete medium containing 100 µg/ml adenine before shifting to 37°C for 4h. Samples were taken every 30’ and further stained for 15’ with 5 µg/ml Hoechst 33258 to visualize the plasma membrane. Since we observed no delay in anaphase for *top2-5* (panels **a** and **b**), we pooled data from 150’-240’ to analyze LDB cells (131 out of 951 cells). The blue arrowhead indicates abscission. Scale bars correspond to 5 µm. BF, bright field. DIC, differential interference contrast.

An unexpected pattern observed in the time course experiments with H2A-GFP was the relative low frequency of chromatin anaphase bridges (i.e., stretched histone-labelled DNA across the bud neck) in both *top2-ts* mutants (Figure 1a, lower panels). Instead, the above-mentioned SDB was the distinctive pattern of *top2-ts* relative to *TOP2*, especially in *top2-4*. In order to shed more light on this phenotype, we filmed cells growing asynchronously and focused on those cells close to reaching anaphase. We observed that all *TOP2* and most *top2-5* cells quickly divide and fully segregate the two nuclear masses to the daughter cell poles (Figure 1b, upper series). By contrast, in the case of *top2-4* we often observed long-lasting histone-labelled SDB cells. Interestingly, this nuclear phenotype was rather dynamic and sometimes became a genuine chromatin bridge during filming (Figure 1b, lower series). To confirm that SDB was a proper split of nuclear masses, we took samples and stained the DNA with DAPI. In all cases, including the SDB phenotype, the DAPI-stained signal overlapped with the H2A-GFP (Figure 1c). These observations indicate that anaphase bridges and SDB are dynamically exchangeable. In the case of *top2-5 HTA2-GFP*, where we observed neither chromatin bridges nor SDBs during the time course and the movies, further labelling with the ultrafine bridge (UFB) marker Dpb11 (Chan, North, and Hickson 2007; Germann et al. 2014), as well as the DNA and plasma membrane reporter Hoechst 33258, showed that there were very few chromatin bridges (8% [4% - 14%]) or UFBs (3% [1% - 8%]) connecting the split nuclear masses in LDB cells. Furthermore, the pre-abscission/post-abscission ratio (1:4) supports the conclusion that most LDB cells have completed cytokinesis (Figure 1d).

Finally, the use of live cell imaging also allowed us to visualize other striking phenotypes that took hours to develop. Although these phenotypes occurred in less than 10% of the cells, they were significant in that they demonstrate the uncoupling of nuclear division and the cell cycle in *top2-ts* (e.g., rebudding before splitting the chromatin bridge) (Figure S1).

### Synthetic genetic array analysis reveals mitotic exit and cytokinesis as deleterious enhancers of transient Top2 inactivation

The prevalence of the dynamic SDB phenotype in *top2-4* relative to *top2-5* raised, in turn, the question of why this difference occurred. In order to search for players that may influence the different behaviours of the *top2* alleles, and to define its biological consequence, we carried out a synthetic genetic array (SGA) analysis with two collections of yeast mutants: the haploid gene deletion collection of non-essential genes (4322 knockout strains) and a collection of ts alleles for essential genes (1231 ts strains) (Li et al. 2011; Tong et al. 2001; Giaever et al. 2002). SGA allows screening for genetic interactions in *Saccharomyces cerevisiae* by comparing the fitness of the colonies of all the different mutant combinations (i.e., single *top2-ts* mutants vs single mutants in the collections vs double *top2-ts*/collection mutants). In addition, we decided to perform the SGA analysis in three different conditions that modify Top2 activity.

In our first analysis, we grew all the strains constantly at the permissive temperature (25ºC), and compared the collection of *top2-4* and *top2-5* double mutants with the corresponding *TOP2* counterparts as references. Thermosensitive alleles are expected to have mild reduced fitness at permissive temperatures and, in our case, we indeed detected genetic interactions at 25 ºC, 139 in *top2-4* and 167 in *top2-5* (Figure 2a, Tables S1 & S2). Eighty-four interactions were shared between *top2-4* and *top2-5*, and these interacted with both alleles in a similar way, either positively or negatively (Figure 2b,Table S3). Among them was *top1Δ*, which is well known to have synthetic sickness with *top2-ts* alleles (Kim and Wang 1989). However, most of the observed genetic interactions were positive, which is typically a sign of suppressive pathways.

**Figure 2.**
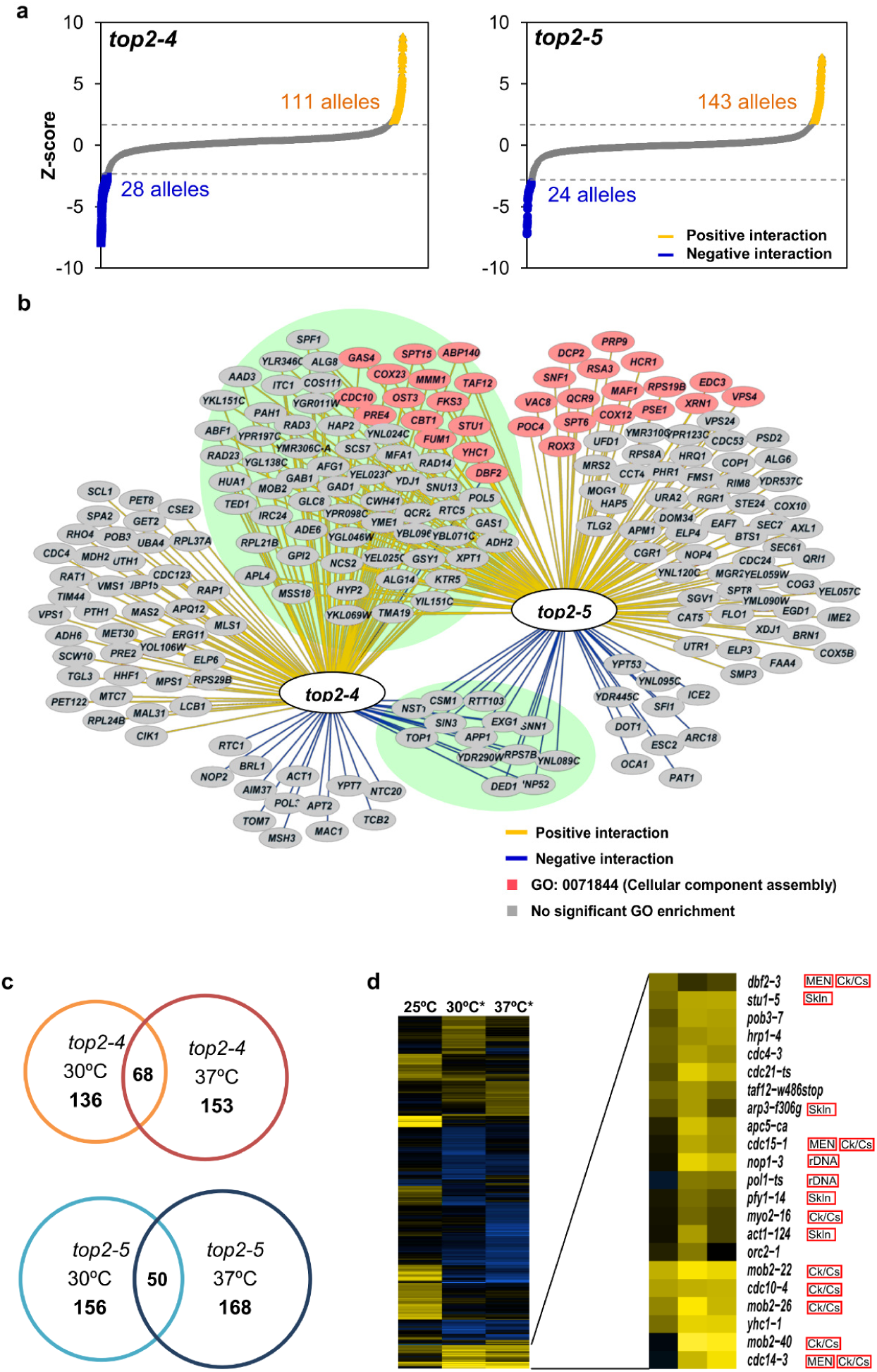
SGA analyses identify the Mitotic Exit Network (MEN) as deleterious enhancers of Top2 downregulation. **(a)** The z-scores of the genetic interactions detected at permissive temperature (steady growth at 25ºC) are plotted. The *top2-ts* arrays of mutants were compared to the *TOP2* arrays as a control. 5553 alleles were screened in total. Positive and negative genetic interactions that meet the cutoffs (>2 or < −2) are indicated. **(b)** Network of the genes that interact with *top2-4* and *top2-5*. Green background encircles the 84 shared genes between the two *top2-ts*. Pink nodes indicate the positive interaction between *top2-5* and genes involved in the aggregation and bonding of cellular components (GO: 0071844). **(c)** The numbers of genetic interactions identified during steady growth at 30ºC and after 6h incubation at 37ºC are shown. Reddish circles, positive interactions; bluish circles, negative interactions. **(d)** Heatmap of *top2-5* SGA scores (those with p<0.05). Red, positive interactions; green, negative interactions; black, no interaction. The 25ºC column represents the z-score obtained comparing *top2-5* versus *TOP2* at 25ºC, whereas the 30ºC and 37ºCx6h columns compare *top2-5* at these temperature regimes versus *top2-5* at 25ºC. MEN, mitotic exit network; Ck/Cs, Cytokinesis/Cell separation; Skln, cytoskeleton; rDNA, ribosomal DNA metabolism.

Next, we sought to study the changes that increasing the temperature would produce in the observed genetic interactions. We opted for two different incubations: one set of arrays was grown at semipermissive temperature (30ºC) for 2 days, and another set was incubated at 37ºC for 6 hours (i.e, enough to complete one cell cycle without Top2 on solid media), and then shifted to 25ºC to allow growth of survivors. In both cases we used the *top2-ts* arrays that were grown at 25ºC as controls to compare the colony size. Thus, we obtained a large number of new interactions that, unlike during growth at 25ºC, were mostly negative (Figure 2c, Tables S4 & S5). Ontological classification of significant interactions revealed common negative interactions at 37ºC × 6h with bioenergetics and autophagy (Table 1). This could be related to the heat shock treatment and putative roles of Top2 during chromosome reshape and transcription reprogramming to cope with this stress (Pommier et al. 2016). Importantly, this classification also spotlighted a high number of positive interactions between *top2-5* grown at 30ºC and/or 37ºC × 6h and thermosensitive alleles related to anaphase/telophase progression (Figure 2d, Table 1). Many of the genes belong to the Mitotic Exit Network (MEN), such as *CDC14* and *CDC15,* whereas others are related to the cytoskeleton or the ribosomal DNA (rDNA) metabolism, which are known to undergo important modifications during anaphase (Machín et al. 2016). To a lesser extent, *top2-4* was also enriched in mitotic division alleles when incubated at 37ºC × 6h, some of which overlap with those of *top2-5* (e.g. *STU1, CDC10* and *MOB2*) (Tables S4 & S5).

**Table 1.**
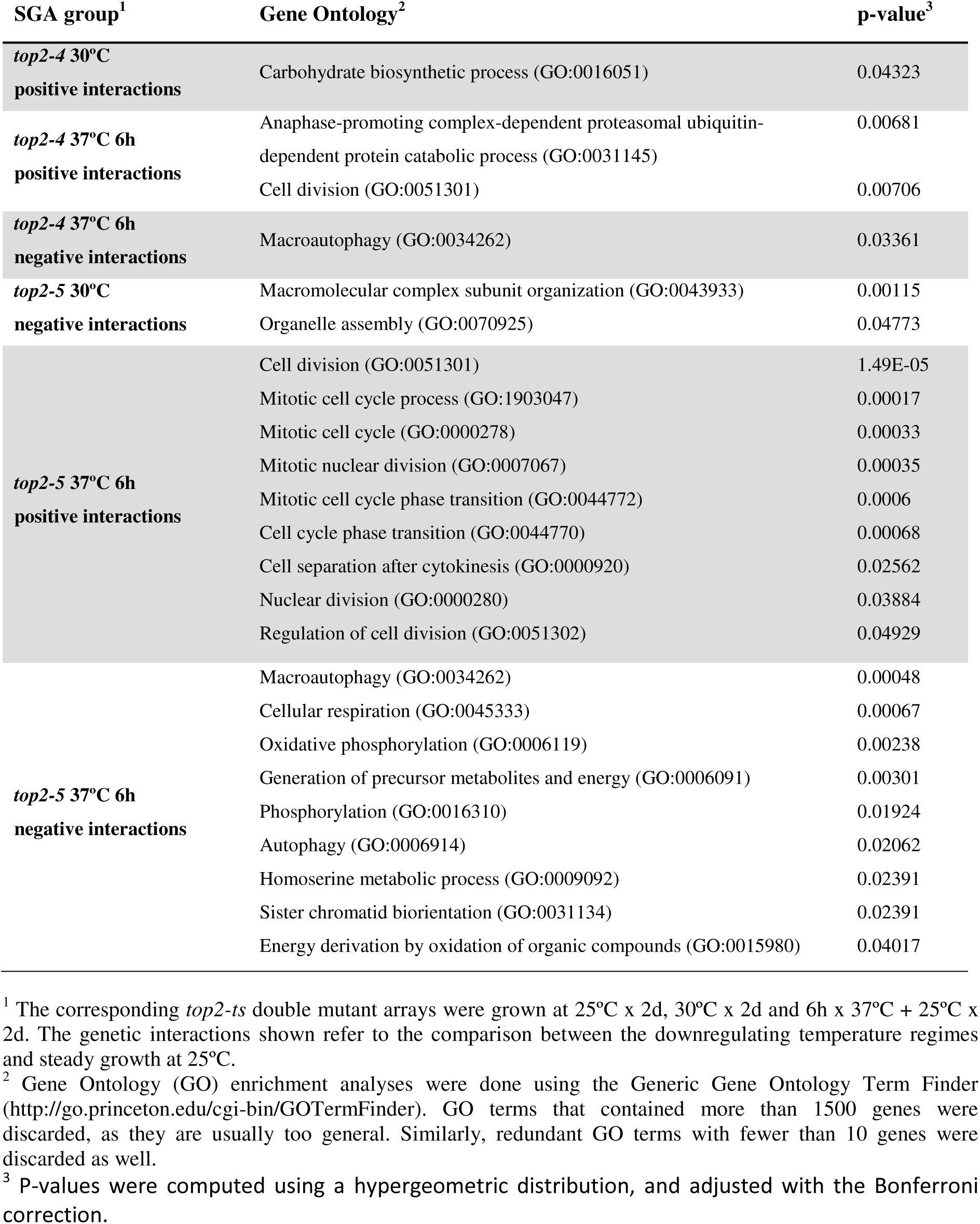
Significant biological processes that genetically interact with the *top2-ts* at different downregulating regimes.

### Anaphase bridges in *top2-ts* are stabilized by preventing mitotic exit

Since MEN and cytokinesis defects were the clearest positive genetic interactions with *top2-ts*, we tested the consequence of disrupting MEN/cytokinesis in the original *top2-ts* strains.

We constructed *top2-5 cdc15-2* and *top2-5 cdc14-1* double ts mutants and arrested these strains in telophase, together with the corresponding *TOP2 cdc15-2/cdc14-1* controls. We found that *top2-5 cdc15-2* could not resolve the histone-labelled anaphase bridge (89% [84%-93%] dumbbells with clearly visible chromatin bridges at 240’; one-third of them having a “short-distance” chromatin bridge phenotype, one-third having a “long-distance” chromatin bridge, and another one-third having a complex string structure) (Figure 3a). By contrast, full segregation (96% [93%-98%]) of the histone signal was seen in all *TOP2 cdc15-2* cells. In the case of *top2-5 cdc14-1*, the nuclear segregation was worse; we observed that most of the nucleus was in the bud (Figure 3b; 84% [77%-90%]). Taking into account that *cdc14-1* strains are known to form an anaphase bridge that comprises the rDNA and also tend to mistakenly segregate the nucleus to the daughter cell (Machín et al. 2005; Machín et al. 2016; Ross and Cohen-Fix 2004), the synergistic nuclear segregation defect in *top2-5 cdc14-1* is not that surprising. Remarkably, this highly asymmetric segregation is the hallmark of mammal epithelial cells treated with Top2 catalytic inhibitors (Gorbsky 1994; Wheatley, O’Connell, and Wang 1998). Overall, these two double mutants demonstrate that the *top2-5* strain gives rise to actual chromatin bridges and that the contraction of the cytokinetic furrow, or an alternative MEN-mediated activity, can quickly split this bridge apart.

**Figure 3.**
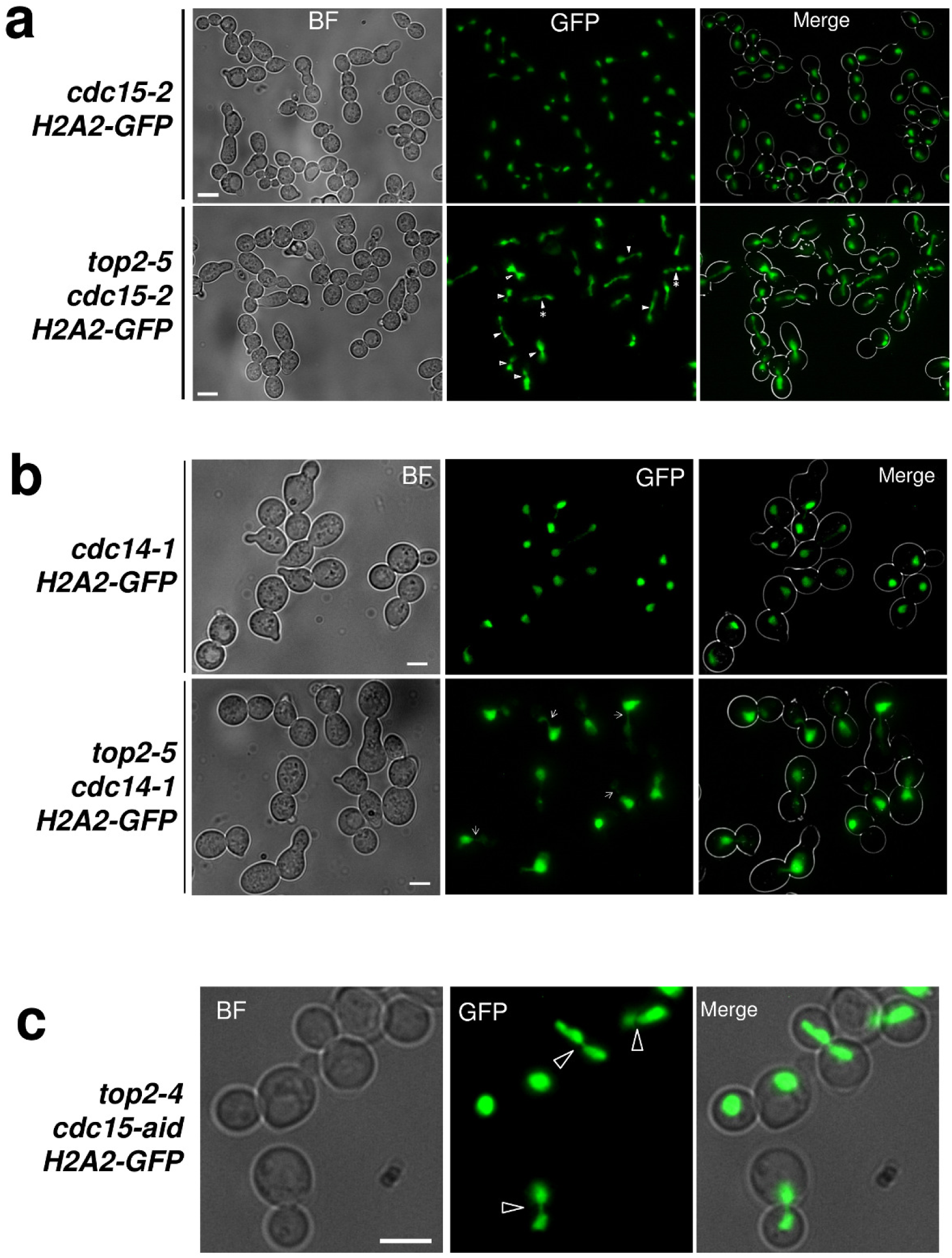
The Mitotic Exit Network (MEN) destabilizes top2-mediated chromatin anaphase bridges. **(a)** The *cdc15-2 HTA2-GFP* and *top2-5 cdc15-2 HTA2-GFP* strains were synchronized in G1 at the permissive temperature (25ºC) and then released into a synchronous cell cycle at 37ºC for 4 h. Samples were taken at the end of the experiment and analysed by fluorescence microscopy. **(b)** The *cdc14-1 HTA2-GFP* and *top2-5 cdc14-1 HTA2-GFP* strains were treated as in panel A. **(c)** The *top2-4 cdc15-aid HTA2-GFP* strain was synchronized in G1 at 25ºC and then released at 37ºC in the presence of 1 mM of the auxin indolacetic acid (IAA) for 3 h. Scale bars correspond to 5 µm. BF, bright field. Hollow arrowheads point to examples of short-distance anaphase bridges. Filled arrowheads point to examples of long-distance anaphase bridges (with an asterisk to highlight the complex strung chromatin bridges). All arrowheads point exactly at the bud neck.

We also made a *top2-4 cdc15-aid* strain. We used *cdc15-aid* (Cdc15 depletion by auxin addition rather than temperature shift) because it was difficult to phenotypically confirm *top2-4 cdc15-2* as *top2-4* alone arrested in telophase (Figure 1a). We also observed chromatin bridges when both Top2-4 and Cdc15-aid were depleted (33% [24%-43%]). In this case, most chromatin bridges had a SDB-like appearance (Figure 3c).

### Conclusions and perspectives

In this work, we have characterized the consequences of depleting yeast cells of Top2 using two *top2-ts* alleles that differ in their resistance to chemotherapeutic Top2 poisons. As previously reported, the first cell cycle takes place with normal kinetics until anaphase, when *top2-ts* form anaphase bridges. We have shown that *top2-5* (poison-resistant) splits apart this bridge more quickly than *top2-4* (poison-sensitive) and is more susceptible to the execution of mitotic exit. Our results point towards upregulation of mitotic exit and/or cytokinesis as new putative targets to synergistically promote cell death upon Top2 downregulation/mutation in cancer cells.

## MATERIALS AND METHODS

### Yeast strain construction, cell cycle experiments and fluorescence microscopy

All the strains used in this work are listed in Table S6 together with their relevant genotypes. C-terminal tagging with GFP/RFP variants or auxin-based degron system, gene deletions and ts allele transfers were made using standard PCR methods as described before (Tong et al. 2001; Janke et al. 2004; Nishimura et al. 2009).

Cell cycle time course experiments and fluorescence microscopy were performed as described before (Quevedo et al. 2012; García-Luis and Machín 2014; Silva et al. 2012). DNA was stained using DAPI (Sigma-Aldrich) at 4 µg/ml final concentration after keeping the cell pellet 24h at −20ºC. Plasma membrane was stained with 5 µg/ml Hoechst 33258 (Sigma-Aldrich) for 15 min at 37°C before processing for fluorescence microscopy. For the time-lapse movies, an asynchronous culture was concentrated by centrifugation to 3 OD_600_ equivalents and plated on YPDA (YPD, agar 2% w/v). Patches were made from this plate and mounted on a microscope slide. They were incubated at 37ºC in high humidity chambers to avoid drying of the agarose patch.

95% confidence intervals for proportions of selected cell phenotypes were calculated assuming a binomial distribution and are indicated in the main text between brackets.

## Synthetic genetic array analyses

Synthetic genetic array (SGA) was performed as described before (Li et al. 2011; Tong et al. 2001). In order to make the strain arrays (described in detail in Figure S2), we first replaced the *TOP2* locus in the haploid *MAT*α strain Y7092 with our query *top2-ts* alleles attached to the selection marker *natMX4* (resistance to nourseothricin). For consistency, we also attached the *natMX4* marker to our reference *TOP2* Y7092 strain. The new *natMX4* strains were then mated with the *MAT***a** mutant colletions (4322 knockout strains for non-essential genes plus 1231 strains with thermosensitive alleles for essential genes.). These panels of *MAT***a** strains bear the *kanMX4* marker (resistance to G418) at the mutated loci. Diploids were selected on YPD plates containing both nourseothricin and G418, and later sporulated and selected for *MAT***a** haploids containing both markers. Once the *TOP2*, *top2-4* and *top2-5* arrays were constructed they were replicated onto plates with the same medium used in the *MAT***a** selection and exposed to the different temperature regimes described in the Results section.

The image analysis, processing and fitness scoring of the arrays were done using SGAtools (http://sgatools.ccbr.utoronto.ca/about) (Wagih et al. 2013). Gene Ontology (GO) enrichment analysis was performed using the Generic Gene Ontology Term Finder (http://go.princeton.edu/cgi-bin/GOTermFinder) of Princeton University (Boyle et al. 2004). Networks were made with Cytoscape v3.3.0 (http://www.cytoscape.org/cy3.html) (Shannon et al. 2003).

## ACKNOWLEDGMENTS

We thank other members of the labs for fruitful discussions and help. This work was supported by the research grants PI12/00280 and BFU2015-63902-R to FM. These grants were funded by the Spanish “Instituto de Salud Carlos III” and the Spanish Ministry of Economy and Competitiveness, respectively. Agencia Canaria de Investigación, Innovación y Sociedad de la Información supported CR through a predoctoral fellowship (TESIS20120109). All these programs were co-financed with the European Commission's ERDF structural funds. The Danish Agency for Science, Technology and Innovation (DFF) and the Villum Foundation supported the work performed by ML. Funding to GWB was provided by the Canadian Cancer Society Research Institute (Impact grant 702310) and the Canadian Institutes of Health Research (grant MOP-79368). The authors declare no conflicts of interest.

